# Polarly localized EccE_1_ is required for ESX-1 function and stabilization of ESX-1 membrane proteins in *Mycobacterium tuberculosis*

**DOI:** 10.1101/820324

**Authors:** Paloma Soler-Arnedo, Claudia Sala, Ming Zhang, Stewart T. Cole, Jérémie Piton

**Author notes:** Correspondence to: Prof. Stewart T. Cole, Institut Pasteur, 25-28 Rue du Dr Roux, 75015 Paris, France. Dr. Jérémie Piton, Global Health Institute, École Polytechnique Fédérale de Lausanne, Station 19, 1015 Lausanne, Switzerland. Ming Zhang, Walter and Eliza Hall Institute of Medical Research, Parkville Victoria, Australia.

## Abstract

*Mycobacterium tuberculosis* is a slow-growing intracellular bacterium with the ability to induce host cell death and persist indefinitely in the human body. This pathogen uses the specialized ESX-1 secretion system to secrete virulence factors and potent immunogenic effectors required for disease progression. ESX-1 is a multi-subunit apparatus with a membrane complex that is predicted to form a pore in the cytoplasmic membrane. In *M. tuberculosis* this complex is composed of five membrane proteins: EccB_1_, EccCa_1_, EccCb_1_, EccD_1_, EccE_1_. In this study, we have characterized the membrane component EccE_1_ and found that deletion of *eccE*_1_ lowers the levels of EccB_1_, EccCa_1_ and EccD_1_ thereby abolishing ESX-1 secretion and attenuating *M. tuberculosis ex vivo*. Surprisingly, secretion of EspB was not affected by loss of EccE_1_. Furthermore, EccE_1_ was found to be a membrane- and cell-wall associated protein that needs the presence of other ESX-1 components to assemble into a stable complex at the poles of *M. tuberculosis*. Overall, this investigation provides new insights into the role of EccE_1_ and its localization in *M. tuberculosis*.

**IMPORTANCE:** Tuberculosis (TB), the world’s leading cause of death of humans from an infectious disease, is caused by the intracellular bacterium *Mycobacterium tuberculosis*. The development of successful strategies to control TB requires better understanding of the complex interactions between the pathogen and human host. We investigated the contribution of EccE_1_, a membrane protein, to the function of the ESX-1 secretion system, the major virulence determinant of *M. tuberculosis*. By combining genetic analysis of selected mutants with eukaryotic cell biology and proteomics, we demonstrate that EccE_1_ is critical for ESX-1 function, secretion of effector proteins and pathogenesis. Our research improves knowledge of the molecular basis of *M. tuberculosis* virulence and enhances our understanding of pathogenesis.

## INTRODUCTION

*Mycobacterium tuberculosis* is a slow-growing intracellular pathogen with the ability to infect and survive inside macrophages. Protein secretion is essential for mycobacterial virulence and host-pathogen interactions (1). The *M. tuberculosis* genome encodes five specialized secretion systems, referred to as ESX or Type VII systems, termed ESX-1 to ESX-5 (2). ESX systems are multi-subunit apparatuses that share similar structure and secrete related proteins but have different functions (3, 4).

ESX-1 is involved in virulence factor secretion and pathogenesis. It is essential for phagosomal rupture thereby allowing translocation to the cytosol where bacteria can induce host cell death and subsequently spread to neighboring cells (6–9). Deletion or inactivation of ESX-1 does not affect growth *in vitro* but causes attenuation of virulence in infection models (9, 10). Indeed, loss of eight genes (RD1) from the *esx-1* locus is the primary attenuating deletion of the live tuberculosis vaccine, *Mycobacterium bovis* BCG (11). The locus encoding the ESX-1 secretion system spans 20 genes and codes for secreted factors and structural components. Additionally, the distal *espACD* operon, encoding three proteins (EspA, EspC and EspD), is also required for full ESX-1 activity (12, 13).

The main ESX-1 substrates, EsxA and EsxB, are major virulence factors and amongst the most potent T-cell antigens of *M. tuberculosis* (14, 4, 15). Besides these two substrates, other proteins are also secreted through the ESX-1 machinery such as EspB, EspA and EspC. Importantly, EsxA/EsxB and EspA/EspC, but not EspB, are co-dependent for secretion (12, 16, 17). Depletion of any of these proteins leads to severe attenuation in cellular and animal models of infection.

ESX-1 has a set of conserved components, with paralogs in all mycobacterial ESX systems, that have been shown to contribute to the ESX secretion process in at least one mycobacterial species (6, 9, 10, 18–23). The five conserved elements EccB_1_, EccCa_1_, EccCb_1_, EccD_1_ and EccE_1_, are likely to be membrane proteins and the core components of the ESX-1 machinery. Paralogs of these proteins encoded by the *esx-5* locus form the membrane complex of the ESX-5 secretion system in *Mycobacterium bovis*, *Mycobacterium marinum* and *Mycobacterium xenopi* (6, 23, 24). New structural insight into the membrane complex indicated that the four components Ecc(BCDE)_5_ are present in equimolar amounts and adopt a hexameric arrangement around a central pore (6, 23, 24). Complex formation of ESX-1 membrane proteins has also been analyzed in *M. marinum*, showing the same composition and size as the ESX-5 complex (23). Considering the conserved nature of the Ecc proteins, the structure of the ESX-1 membrane complex should resemble that of ESX-5.

Another conserved component of the ESX-1 apparatus is MycP_1_, a membrane protein, which is not part of the core membrane complex but loosely associated with it (23). MycP_1_ is a subtilisin-like serine protease that cleaves EspB in the periplasmic space (23). Besides its role in substrate processing, MycP_1_ plays a second role in the secretion process by stabilizing the ESX-1 membrane complex (23). In the *esx-1* locus of *M. tuberculosis, mycP*_*1*_ is situated upstream of *eccE*_*1*_ and the two genes are co-transcribed (25). EccE_1_ is the conserved element that has been the least explored, especially in *M. tuberculosis* where no experimental work has been reported yet. In *M. smegmatis*, the homologue of EccE_1_ is required for EsxA and EsxB secretion (19). Previous work on the composition and structure of the ESX-5 membrane complex demonstrated that EccE_5_ is located at the perimeter of the membrane complex and is important for its formation and stability (6, 24).

Although the molecular mechanisms underlying the ESX secretion process are not fully understood, studying individual components has provided important insight. Localizing active ESX systems can also offer valuable information about the mechanism of secretion. ESX-1-related proteins have been visualized at the cell poles of *M. marinum* and *M. smegmatis* (26, 27). Nonetheless, information about the localization of the entire ESX-1 system in *M. tuberculosis* is missing.

In this study, we explored the role of EccE_1_ in *M. tuberculosis*. We show that EccE_1_ is required for *ex vivo* virulence, for stabilizing ESX-1 membrane proteins and for secretion of EsxA, EsxB, EspA and EspC. Moreover, we investigated the localization of the protein and its incorporation into a functional ESX-1 system. Our findings indicate that EccE_1_ is a membrane-protein that requires other ESX-1 components to form a stable complex at the poles of *M. tuberculosis*.

## RESULTS

### Mutant construction

To assess the role of EccE_1_ in ESX-1-related functions, we first constructed an *eccE*_*1*_ deletion mutant in *M. tuberculosis*. As *eccE*_*1*_ is in the same transcriptional unit as *mycP*_*1*_ (25), we initially deleted the entire *mycP*_*1*_-*eccE*_*1*_ region from the chromosome by allelic exchange. Whole genome sequencing confirmed the complete deletion of the *mycP*_*1*_-*eccE*_*1*_ coding sequences (CDS) and the absence of other mutations in the genome. The resulting *mycP*_*1*_-*eccE*_*1*_ double mutant was then partially or fully complemented with an integrative plasmid bearing either the single genes *mycP*_*1*_ or *eccE_1_*, or the entire *mycP*_*1*_-*eccE*_*1*_ locus under the control of the PTR promoter. As a result, we obtained four strains: a double mutant ∆*mycP*_*1*_-*eccE*_*1*_, a fully complemented strain ∆*mycP*_*1*_-*eccE*_*1*_/*mycP*_*1*_-*eccE*_*1*_ and two partially complemented strains ∆*mycP*_*1*_-*eccE*_*1*_/*mycP*_*1*_ and ∆*mycP*_*1*_-*eccE*_*1*_/*eccE*_*1*_ which correspond to the single mutants ∆*eccE*_*1*_ and ∆*mycP*_*1*_, respectively.

The four strains were subsequently analyzed by RNA sequencing and the transcriptional profile compared to that of the H37Rv wild type (WT) strain. As anticipated, no de-regulated genes were found except for the deleted genes, suggesting that the excision of *eccE*_*1*_-*mycP*_*1*_ and their complementation *in trans* did not impact the transcription of other genes. In the complemented derivatives, the expression levels of *mycP*_*1*_ and *eccE*_*1*_ were very similar to those measured in the WT strain (Table S1).

The ability of the mutant strains to grow in synthetic media was explored. The Δ*eccE*_1_ mutant, as well as the ΔmycP_1_ and the Δ*mycP*_1_-*eccE*_*1*_ strains, exhibited growth kinetics similar to that of the WT strain (Fig S1). These results indicate that *eccE*_*1*_ and *mycP*_1_ are not required for *in vitro* growth of *M. tuberculosis*.

### EccE_1_ and MycP_1_ are not involved in susceptibility to drugs targeting the cell envelope

Previous studies on the ESX-5 system demonstrated that disruption of this apparatus in *M. tuberculosis* reduces cell wall integrity (18). Consequently, susceptibility to various antibiotics targeting several steps in the cell wall biosynthetic pathway greatly increased (18). To test whether its paralog, the ESX-1 system, is also involved in maintaining the stability of the cell envelope, the drug susceptibility of the mutants to various cell wall targeting antibiotics was evaluated using the resazurin-based microdilution assay (REMA) (28). The Δ*eccE*_1_ mutant and the WT strain displayed similar minimal inhibitory concentrations (MIC) for all antibiotics tested, as did the Δ*mycP*_1_ and the Δ*mycP*_1_-*eccE*_*1*_ mutants (Table S2). We concluded that the absence of either *eccE*_1_ or *mycP*_1_ did not impact susceptibility to β-lactams (penicillins and cephalosporins) or to glycopeptides (vancomycin). In contrast to ESX-5, a secretion system previously described as essential for cell wall integrity in *M. tuberculosis*, ESX-1 inactivation did not impact susceptibility to the antibiotics tested.

### EccE_1_ and MycP_1_ are required for *M. tuberculosis*-mediated cell death

EccE was identified as a peripheral component of the ESX membrane complex, but further information about its role in *M. tuberculosis* is missing. To investigate the implication of EccE_1_ in ESX-1-mediated function, we exploited the ability of *M. tuberculosis* to induce host cell death, which is linked to ESX-1-related secretion (5, 7). To this aim, THP-1 human macrophages were infected with the *mycP*_*1*_ and *eccE*_*1*_ mutants as well as with the WT strain and the survival of macrophages was monitored at 72h post-infection. In the presence of WT *M. tuberculosis*, the survival of macrophages significantly decreased compared to the uninfected control. In contrast, when macrophages were infected with strains lacking either *eccE*_*1*_ and/or *mycP*_*1*_, the macrophages survived to levels similar to the non-infected control. Importantly, the complemented mutant restored the WT phenotype (Fig 1). This result indicates that *M. tuberculosis* needs both EccE_1_ and MycP_1_ for cytotoxicity and that the absence of either of these ESX-1 components leads to complete attenuation of *M. tuberculosis ex vivo*.

**FIG 1.**
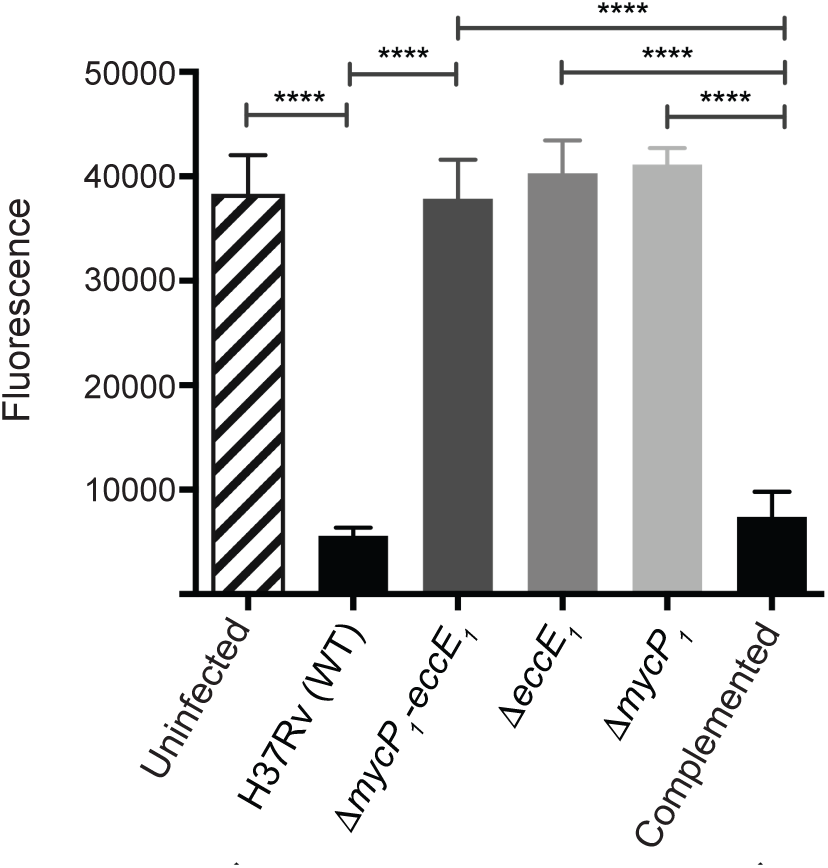
EccE_1_ and MycP_1_ are required to induce *M. tuberculosis*-mediated cell death. Cell viability of THP-1 human macrophages at 72h post-infection. Uninfected cells were used as a control for macrophage survival and the H37Rv WT strain as a positive control for *M. tuberculosis*-mediated cell-death. The Δ*eccE*_1_, Δ*mycP*_1_-*eccE*_1_ and Δ*mycP*_1_ mutants did not affect survival of THP-1 macrophages in contrast to the complemented strain. Columns represent mean values from three independent biological replicates and error bars show the standard deviation. One-way ANOVA followed by Tukey’s multiple comparison test was used for statistical comparison. ****P-value <0.001.

To exclude an infection defect that could impact macrophage survival and to validate the attenuated phenotype displayed in the absence of *eccE*_*1*_ and/or *mycP1*, we determined the levels of intracellular bacteria at 4h post-infection by colony forming units (CFU) enumeration. The bacterial burden was similar for the WT and the mutants, indicating that all strains were phagocytosed with similar efficiency (S2 Fig). These results demonstrated that *M. tuberculosis* does not require EccE_1_ or MycP_1_ to infect macrophages, but it does need both components to induce host cell death.

### EccE_1_ and MycP_1_ are essential for ESX-1-protein-secretion

To determine if EccE_1_ and MycP_1_ are crucial to the function of the ESX-1 machinery, we checked whether the deletion of *eccE*_*1*_ and/or *mycP*_*1*_ impacts ESX-protein secretion. To this end, cell lysates and culture filtrates of WT and mutant bacteria were prepared and analyzed by immunoblotting. The presence of Ag85B, an ESX-1 independent secreted protein, and the absence of the cytosolic GroEL2 protein in the culture filtrates served as controls. As expected, EsxA and EsxB were present in the secretome of the WT *M. tuberculosis* strain. However, EsxA and EsxB were not detected in culture filtrates of strains lacking *eccE*_*1*_ and/or *mycP*_*1*_ although both substrates were present in the cell lysates. Furthermore, secretion of EsxA and EsxB was restored in the fully complemented strain Δ*mycP*_1_-*EccE*_1_/*mycP*_1_-*EccE*_*1*_ (Fig 2A). These data indicate that both EccE_1_ and MycP_1_ are required for secretion of the two main ESX-1 substrates. In contrast, EspB, another known ESX-1 substrate, was found in the cell lysates and culture filtrates of the WT, mutant and complemented strains (Fig 2A), suggesting that secretion of EspB is independent of EccE_1_ or MycP_1_. Overall, we conclude that expression of EsxA, EsxB and EspB is not affected by the absence of EccE_1_ or MycP_1_. However, both ESX-1 membrane proteins are essential for EsxA and EsxB secretion but dispensable for the release of EspB into the culture filtrate.

**FIG 2.**
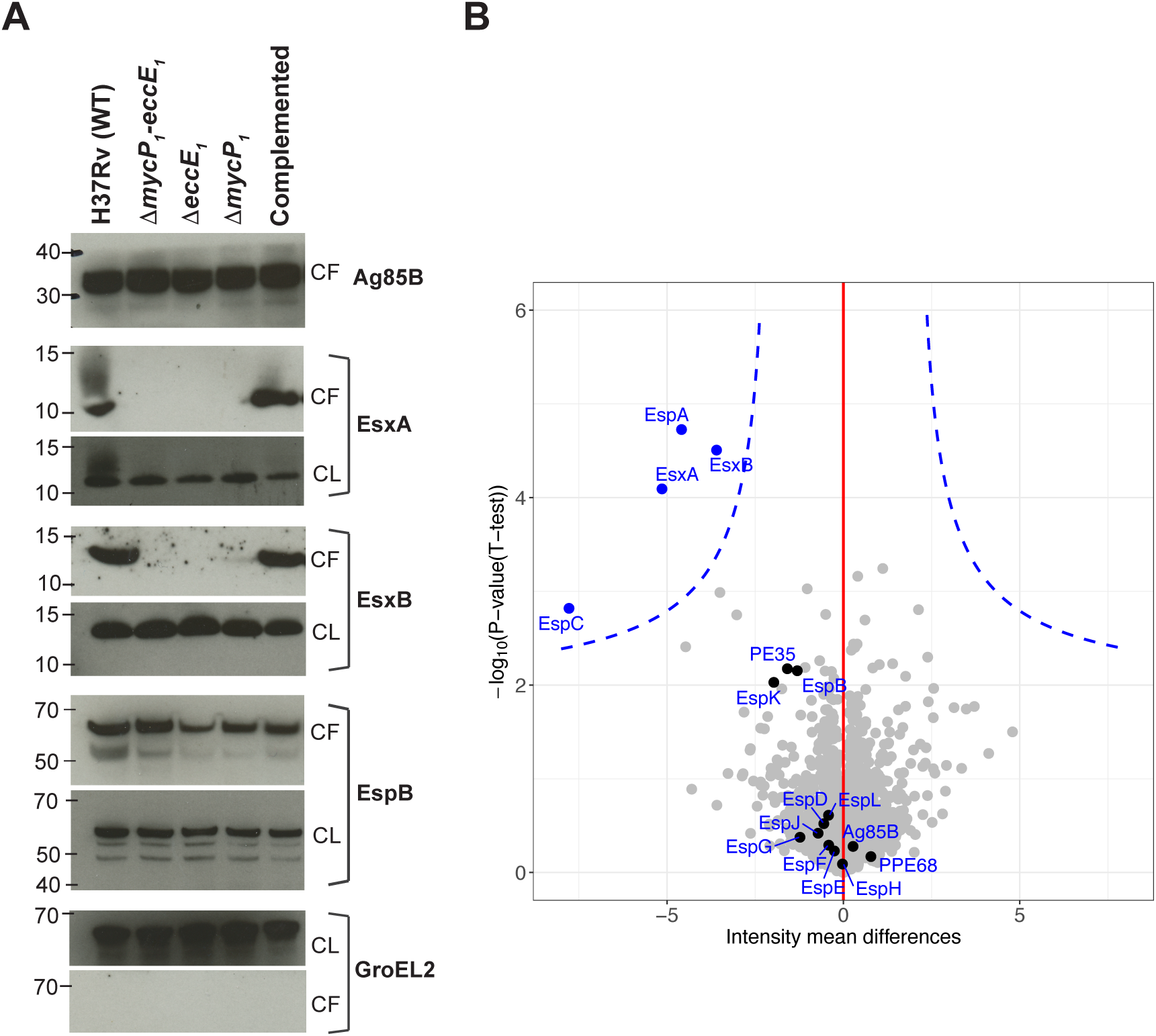
EccE_1_ and MycP_1_ are required for secretion of EsxA, EsxB, EspA and EspC but not EspB. **A.** Immunoblots of culture filtrates (CF, 15 μg per well) and whole cell lysates (CL, 10 μg per well). Detection of Ag85B was used as a loading control in the CF and GroEL2 as a loading control in the CL. Secretion of EsxA and EsxB but not EspB is disrupted in the Δ*eccE*_1_, Δ*mycP*_1_-*eccE*_1_, and Δ*mycP*_1_ mutants. **B.** Proteomic analysis of the secreted fraction comparing the Δ*mycP*_1_-*eccE*_1_ mutant with the H37Rv WT strain. Each dot corresponds to an identified protein. Proteins are represented using a volcano plot-based strategy where t test p-values on the -log base 10 scale are combined with ratio information on the log base 2 scale. The blue dotted line represents the significance curve (SO value of 0.5) and it delineates the differentially quantified proteins. Three independent biological replicates per strain were analyzed and a t-test (significance level of 0.05) was used for statistical comparison. The abundance of EsxA, EsxB, EspA and EspC is highly affected in the double mutant Δ*mycP*_1_-*eccE*_1_ compared to the WT strain. EsxA, EsxB, EspA and EspC are the only significantly reduced proteins.

To obtain deeper knowledge into the impact of EccE_1_ and MycP_1_ on *M. tuberculosis* secretion and to identify potential EccE_1_-dependent substrates, the whole secretome of the mutants was analyzed and compared to that of the WT strain (Table S5). Bacteria were grown in the same conditions as previously described for immunoblotting and the secreted fraction analyzed by mass spectrometry. We found that EsxA, EsxB, EspA and EspC were the only proteins significantly reduced in the culture filtrates (Fig 2B). Moreover, in the WT strain, EsxA was in the top ten most abundant proteins. On the other hand, the secretion of this protein greatly decreased (Log_2_ ratio −5.1) when EccE_1_ and MycP_1_ were missing (Table S5). The same trend was observed for EsxB, EspA and EspC whose secretion was also greatly decreased (Log_2_ ratio [-7.8, −3.6]) in the deletion mutants compared to the WT strain (Table S5). It was noteworthy that the level of the other ESX-1-related proteins, including EspB, was not different in the absence or presence of EccE_1_ and MycP_1_ (Fig 2B). Proteins using other secretion systems, such as Ag85B, were secreted at similar levels in the WT and in the mutants (Fig 2B). Taken together, the secretome analysis demonstrated that only secretion of EsxA, EsxB, EspA and EspC is dependent on EccE_1_ and MycP_1_. This analysis also confirmed the previous results obtained by immunoblotting.

### EccE_1_, but not MycP_1_, impacts the stability of various ESX-1 membrane proteins

In order to evaluate whether the absence of EccE_1_ and MycP_1_ could affect the stability of other ESX-1 membrane proteins, we analyzed the intracellular proteome of the mutants (Δ*eccE*_1_, Δ*mycP*_1_, Δ*mycP*_*1*_-*eccE*_*1*_) and compared it to that of the WT strain (Table S6). As expected, EccE_1_ and MycP_1_ were present in the WT and absent in the corresponding mutants. In addition, the intracellular control proteins GroEL2 and RpoB were found at similar levels in all strains. No significant differences were observed for EsxA, EsxB or EspB (Fig 3, S6 Table), thus corroborating the data obtained by immunoblot (Fig 2A). However, in the absence of EccE_1_, the levels of EccCa_1_, EccB_1_ and EccD_1_ were reduced (Figs 3A and 3C, S6 Table). Interestingly, the abundance of these membrane proteins did not change in the Δ*mycP*_1_ mutant compared to the WT strain (Fig 3B, S6 Table). Considering the RNA-seq data, which showed no deregulation of *eccCa*_*1*_, *eccB*_*1*_ and *eccD*_*1*_ transcription (S1 Table), these data suggest that the absence of EccE_1_, but not MycP_1_, impacted the stability of EccCa_1_, EccB_1_ and EccD_1_.

**FIG 3.**
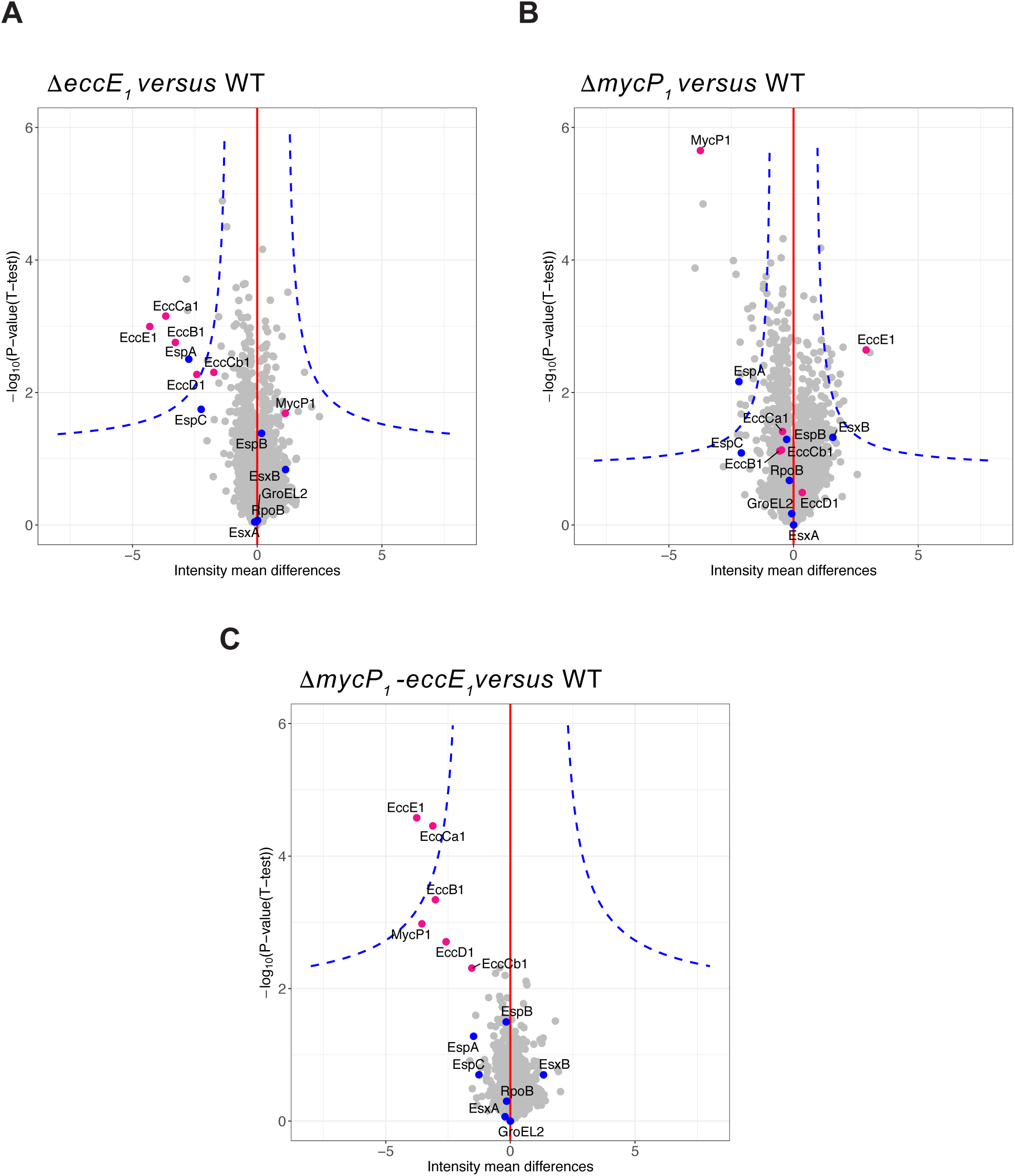
Deletion of EccE_1_, but not MycP_1_, impacts stability of various ESX-1 membrane proteins. Proteomic analysis of the cell lysate fractions comparing different *M. tuberculosis* mutants to the WT strain. Each dot corresponds to an identified protein. Proteins are represented using a volcano plot-based strategy where t test p-values on the -log base 10 scale are combined with ratio information on the log base 2 scale. The blue dotted line represents the significance curve (SO value of 0.5) and it delineates the differentially quantified proteins. Three independent biological replicates per strain were analyzed and a t-test (significance level of 0.05) was used for statistical comparison. In pink: ESX-1 membrane proteins. In blue: ESX-1 secreted proteins and intracellular protein controls. **A**. Δ*mycP*_1_-*eccE*_1_/*mycP*_1_ versus WT. **B**. Δ*mycP*_1_-*eccE*_1_/*eccE*_1_ versus WT. **C**. Δ*mycP*_1_-*eccE*_1_ versus WT.

### EccE_1_ is a membrane and cell wall-associated protein

Since antibodies against EccE_1_ of *M. tuberculosis* were not available, an HA tag was inserted in the C-terminal part of EccE_1_ by genetic engineering, as recently applied to other ESX-1-related proteins with success (29). For this purpose, the double mutant Δ*mycP*_1_-*eccE*_*1*_ was complemented with an integrative plasmid carrying *mycP*_*1*_ and *eccE*_1_-*HA* (Δ*mycP*_1_-*eccE*_1_/*mycP*_1_-*eccE*_1_*HA*). Both genes were present in single copy and expressed under the control of the same promoter, PTR. To ensure that the tag did not interfere with the activity of EccE_1_, the ability of the HA-tagged complemented mutant to induce cell lysis was tested. The THP-1 *ex vivo* model of infection confirmed that the EccE_1_-HA expressing mutant was as cytolytic as the WT strain, thereby implying the presence of a functional ESX-1 system and so, a functional EccE_1_ protein (Fig 4A).

**FIG 4.**
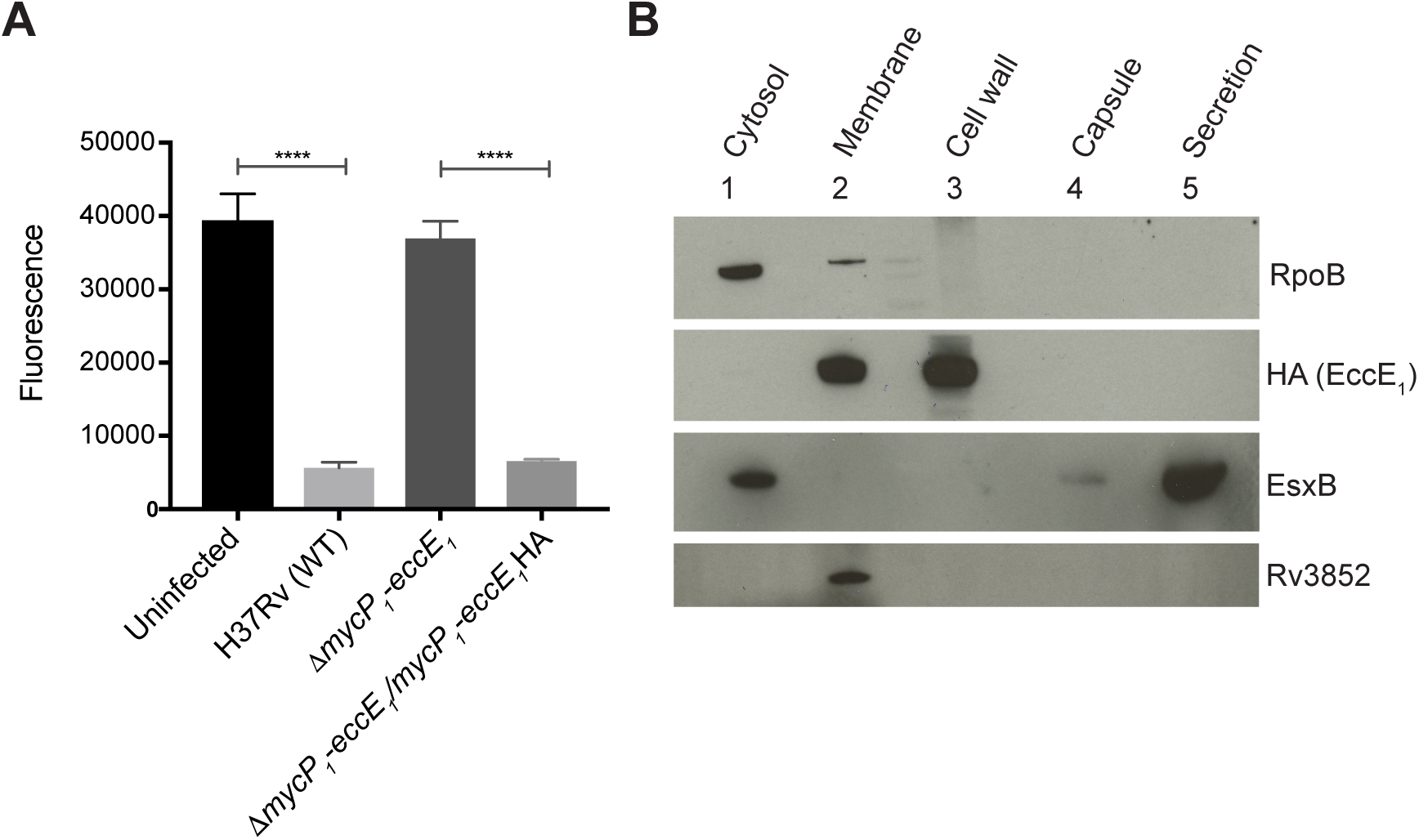
EccE_1_ localizes to the membrane and cell wall fractions of *M. tuberculosis*. **A.** THP-1 macrophage survival at 72h post-infection. Δ*mycP*_*1*_-*eccE*_*1*_/*mycP*_*1*_-*eccE*_*1*_*HA* mutant induces cell-death *ex vivo* indicating the presence of a functional ESX-1 system and therefore that the HA tag at the C-terminal part of the protein did not interfere with EccE_1_ function. Uninfected cells were used as a control for macrophage survival, the H37Rv WT strain and the Δ*mycP*_*1*_-*eccE*_*1*_ mutant served as positive and negative controls, respectively, for *M. tuberculosis*-mediated cell-death. Columns represent mean values from three independent replicates and error bars show the standard deviation. One-way ANOVA followed by Tukey’s multiple comparison test was used for statistical comparison. **** P-value <0.001. **B.** Subcellular fractionation of the Δ*mycP*_*1*_-*eccE*_*1*_/*mycP*_*1*_-*eccE*_*1*_*HA* mutant followed by immunoblotting of each fraction (20 μg per well). Detection of RpoB represented a control for lysis, EsxB for secretion and Rv3852 for the membrane fraction. Anti-HA antibodies localized EccE_1_.

EccE_1_ is predicted to be a membrane protein in *M. tuberculosis*. Subcellular fractionation of the mutant carrying *eccE*_*1*_-*HA* was therefore performed and the cytosolic, membrane, cell wall, capsular and secreted fractions probed by immunoblotting using control proteins of known subcellular location to validate the procedure. RpoB, an RNA polymerase subunit, was found mainly in the cytosol and partly in the membrane. Rv3852, a membrane protein, was only present in the membrane fraction as previously described (30). EsxB, a substrate of the ESX-1 system, was detected in the cytosol and to a lesser extent in the capsule although it was mainly ^found in the culture filtrate. Finally, EccE_1_ was identified in the membrane and cell wall but^ not in other compartments of *M. tuberculosis* (Fig 4B).

### EccE_1_ assembles with other ESX-1 proteins at the poles of *M. tuberculosis*

Fluorescent-fusion proteins have been successfully employed to study ESX-1-related proteins in *M. smegmatis* (27). Overexpression of reporter proteins is usually required for visualization, however, production of a large amount of protein can eventually lead to aberrant phenotypes or artefacts. To avoid this potential problem, we constructed a fluorescent-fusion protein of EccE_1_ with mNeon, a 27 kDa monomeric fluorescent protein reported to be three to five times brighter than GFP and export-competent (31–33). To generate the EccE_1_-mNeon expressing strain, the same strategy used for generating the Δ*mycP*_1_-*eccE*_1_/*mycP*_1_-*eccE*_1_-*HA* was followed. The complemented derivative (Δ*mycP*_1_-*eccE*_1_/*mycP*_1_-*eccE*_1_mNeon) has a single copy of *mycP*_1_ and *eccE*_1_-mNeon genes under the control of the PTR promoter, which provides expression of EccE_1_-mNeon close to physiological levels. To check whether fusion to mNeon impacted EccE_1_ function, the cytotoxic phenotype of the complemented mutant was analyzed using THP-1 macrophages. The EccE_1_-mNeon expressing mutant induced cell death similarly to the WT strain (Fig 5B), suggesting that this mutant can form a functional ESX-1 secretion system. Since the WT copy of EccE_1_ is not present in the mutant and this protein is required for ESX-1 function in *M. tuberculosis*, all active ESX-1 apparatuses in the fluorescent mutant should contain EccE_1_-mNeon proteins.

**FIG 5.**
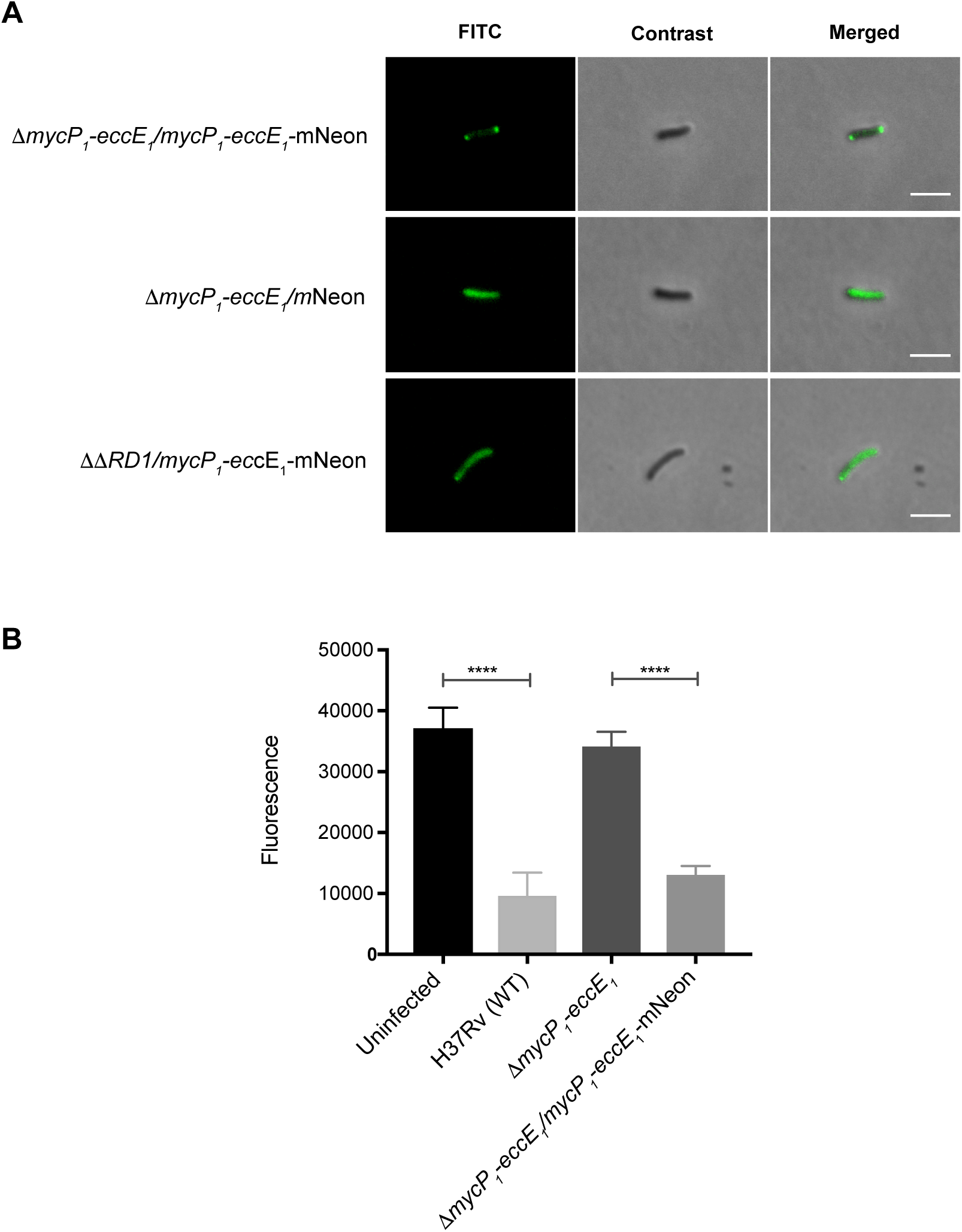
EccE_1_-mNeon and the functional ESX-1 system localize to the poles of *M. tuberculosis*. **A.** EccE_1_-mNeon expressing cells (Δ*mycP*_1_-*eccE*_1_/*mycP*_1_-*eccE*_1_-mNeon) display a fluorescent signal at the poles of the bacilli. When mNeon is expressed without EccE_1_ (Δ*mycP*_1_-*eccE*_1_/mNeon), the fluorescent signal distributes all over the bacterium. Expressing EccE_1_-mNeon in the ΔΔRD1 mutant, which lacks most of the ESX-1 genes, did not result in polar localization, showing that EccE_1_ requires other ESX-1 protein to localize at the poles. The white bar on the right panels represents a 3 μm scale bar. **B.** THP-1 survival at 48h post-infection. The Δ*mycP*_1_-*eccE*_1_/*mycP*_1_-*eccE*_1_-mNeon mutant induces cell-death *ex vivo* suggesting the presence of functional ESX-1 systems and therefore indicating that the addition of mNeon did not interfere with EccE_1_ function. Uninfected cells were used as a control for macrophage survival, the H37Rv WT strain and the Δ*mycP*_1_-*eccE*_1_ mutant were used as positive and negative controls respectively for *M. tuberculosi*s-mediated cell-death. Columns represent mean values from three independent replicates and error bars show the standard deviation. One-way ANOVA followed by Tukey’s multiple comparison test was used for statistical comparison. **** P-value <0.001.

To gain more insight into the localization of EccE_1_ and of the ESX-1 system in *M. tuberculosis*, the fluorescent mutant Δ*mycP*_1_-*eccE*_*1*_/*mycP*_1_-*eccE*_1_mNeon was used to image live bacteria. The double mutant ΔmycP_1_-*eccE*_*1*_ expressing mNeon alone (Δ*mycP*_1_-*eccE*_*1*_/mNeon) and the H37Rv ΔΔRD_1_ mutant (lacking the extended *esx-1* locus) (34) expressing EccE_1_-mNeon (ΔΔRD_1_/*mycP*_*1*_-*eccE*_*1*_mNeon) were used as controls. We observed bi-polar foci in all Δ*mycP*_*1*_-*eccE*_1_/*mycP*_1_-*eccE*_1_mNeon cells examined (S3A Fig). However, cells expressing mNeon alone displayed a diffuse signal throughout the bacterium (Fig 5A), as did the ΔΔRD_1_ mutant expressing EccE_1_-mNeon (Fig 5A and S3B Fig). These results indicate that most of the EccE_1_ proteins localize to the poles of *M. tuberculosis* in the presence of a functional ESX-1 system and suggest that the bi-polar localization may not only correspond to the EccE_1_ protein but to the assembled ESX-1 complex.

## DISCUSSION

In the present study, we constructed and employed mutants and genetic tools to understand the localization and functional role in *M. tuberculosis* of EccE_1_, an as yet unexplored ESX-1 membrane protein. We demonstrate that the absence of EccE_1_ does not impact antibiotic susceptibility or bacterial growth *in vitro* (Fig S1) whereas EccE_1_ is essential for macrophage lysis, a function required for *M. tuberculosis* pathogenesis (Fig 1). To the best of our knowledge, this is the first time that disruption of the core component EccE_1_ in *M. tuberculosis* has proved to lead to complete attenuation *ex vivo*.

Analysis of the ESX-1-secretome explains this phenotype as secretion of EsxA, EsxB, EspA and EspC, which is essential to promote virulence (12, 13, 20), was found to be strongly dependent on EccE_1_ (Fig. 2). Proteomic data showed no significant variations in the intracellular levels of EsxA, EsxB, EspA and EspC when EccE_1_ and MycP_1_ were missing (Fig 3), indicating that the reduced secretion of these ESX-1 substrates is due to disruption of the secretion process and not to decreased protein production. The same phenotype was also observed in the Δ*mycP*_1_ mutant, confirming that MycP_1_ is essential for EsxA/B secretion and virulence in *M. tuberculosis*, as previously reported (21). However, secretion of EspB was not affected by the absence of EccE_1_ and/or MycP_1_ suggesting that secretion of this protein is not ESX-1 dependent (Fig 2) and that another mechanism may be involved, as postulated earlier for EspD (35). In contrast with our findings, EspB secretion was previously reported to require MycP_1_ in both *M. tuberculosis* and *M. marinum* (21, 23). In our investigation, secretion of EspB was analyzed in the *M. tuberculosis* H37Rv strain whereas in previous studies an Erdman background was used (21). However, EspB was also secreted by an attenuated H37Rv ESX-1 mutant lacking EspL while secretion of the main ESX-1 substrates, EsxA, EsxB, EspA and EspC, was significantly reduced (36). Strain differences may thus explain this discrepancy in EspB secretion.

A further strain-dependent difference occurs in the EspB secretion pattern. While EspB is secreted mainly as a cleaved form by the *M. tuberculosis* Erdman strain (21) (Fig S4), both cleaved and full-length EspB are secreted in the same proportions by *M. tuberculosis* H37Rv (Fig S4). This indicates that the genetic background of the *M. tuberculosis* strains should be considered when analyzing EspB secretion.

The analysis of the intracellular proteome also revealed the impact of EccE_1_ on the stability of ESX-1 membrane proteins and, therefore, formation of the ESX-1 membrane complex (Fig 3). It is plausible that in the absence of EccE_1_, the EccB_1_, EccCa_1_, EccD_1_ and EccCb_1_ proteins, may not properly assemble or fold and hence become more vulnerable to degradation. In line with these results, Bosserman and colleagues (37) recently demonstrated that the lack of EccCb_1_ in *M. marinum* led to reduced levels of various ESX-1 membrane proteins including EccE_1_. Together, these results support the hypothesis that loss of a single Ecc protein destabilizes the ESX-1 complex. In contrast, the absence of MycP_1_ did not affect the intracellular levels of the ESX-1 membrane proteins suggesting that the membrane complex is probably formed in the protease-deficient mutant. Consistent with this, previous work on *M. marinum* reported the formation of the ESX-1 core membrane complex in the absence of the MycP_1_ mycosin (23), which is not itself an integral part of the ESX-1 apparatus but nonetheless crucial for its integrity and function (23).

We confirmed that EccE_1_, with its two N-terminal transmembrane regions (38), is indeed a membrane-anchored protein (Fig 4) with its hydrophilic C-terminal part predicted to be in the periplasm. However, detection of the EccE_1_-mNeon protein in the cell wall fraction (Fig 4) as well suggests that EccE_1_ also localizes there and thus behaves similarly to EccB_1_, another core component of ESX-1 with two localizations. Indeed, EccB_1_, a periplasmic protein with a single transmembrane region, was found in both the plasma membrane and cell wall fractions (39). These observations are consistent with the structure of the ESX-5 membrane complex where the soluble domain of EccE_5_ was predicted to be located in the periplasmic space (24).

Two previous studies demonstrated the presence of ESX-1 core components at a single cell pole in *M. marinum* and *M. smegmatis* using immunofluorescence and microscopy analysis of single cells overexpressing fluorescent fusion proteins (26, 27). In this study, we localized EccE_1_-mNeon at both poles of *M. tuberculosis*, using low level expression (Fig 5A), but only when a functional ESX-1 apparatus was present. In its absence, the EccE_1_-mNeon signal was diffuse and observed throughout the cell (Fig 5). These observations suggest that since EccE_1_ is localized at the poles in *M. tuberculosis* the ESX-1 apparatus may be there too. To test this, the EccE_1_-mNeon protein constructed here may prove to be a useful tool for visualizing the nanomachine *in situ* by correlative light-electron microscopy.

## MATERIAL AND METHODS

### Bacterial culture conditions

*M. tuberculosis* was routinely grown in 7H9 broth (supplemented with 0.2% glycerol, 10% ADC, 0.05% Tween-80) or on 7H10 agar (supplemented with 0.5% glycerol and 10% OADC). *Escherichia coli* TOP10 or chemically-competent *E. coli* DH5α cells were used for cloning and plasmid propagation and were grown on LB broth or agar.

### Mutant construction

Deletion of the *mycP*_*1*_-*eccE*_*1*_ region was accomplished in the H37Rv strain by two-step homologous recombination using the pJG1100-derived vector (40). Two fragments of ~900 bp corresponding to the up- and downstream regions of *mycP*_*1*_-*eccE*_*1*_ were PCR-amplified and inserted in pJG1110 vector. The first recombination event was selected on 7H10 plates with hygromycin (50 μg/ml) and kanamycin (20μg/ml). Positive colonies identified by PCR were grown in 7H9 medium with no antibiotics and subjected to a second selection on 7H10 plates supplemented with 2.5% sucrose. The absence of *mycP*_*1*_-*eccE*_*1*_ in the resulting clones was scored by PCR and further confirmed by Southern blotting.

### DNA extraction and whole-genome sequencing

The *M. tuberculosis* Δ*mycP*_1_-*eccE*_1_ mutant was grown in 7H9 broth to OD_600_ 0.8. Bacteria were harvested by centrifugation and DNA was extracted using the QIAmp UCP pathogen kit (Qiagen) as described previously (41). Ilumina libraries were prepared using the Kapa Hyper prep kit as described (41) and quantified using Qubit dsDNA BR Assay Kit (Thermo Fisher Sc). Fragment size was assessed on a Fragment Analyzer (Advanced Analytical Technologies). Finally, libraries were multiplexed and sequenced as 100 base-long single-end reads on an Illumina HiSeq 2500 instrument. Reads were adapter- and quality-trimmed with Trimmomatic v0.33 (42) and mapped onto the *M. tuberculosis* H37Rv reference genome (RefSeq NC_000962.3) using Bowtie2 v2.2.5 (Langmead & Salzberg, 2012).

### Complemented derivative constructions

The integrative pGA44 vector (43) was used to construct all complemented derivates. All plasmids used in this study are listed in Table S3. Briefly, the genes of interest were PCR amplified from *M. tuberculosis* H37Rv genomic DNA (Table S4) and cloned in-frame under the PTR promoter. The monomeric Yellow-Green fluorescent protein, mNeonGreen (32), and the HA tag sequences were cloned at the c-terminal part of *eccE*_1_. All plasmids were checked by Sanger sequencing and were further transformed into competent *M. tuberculosis* Δ*mycP*_1_-*eccE*_1_ cells together with pGA80 (43) which provided the integrase. All strains, plasmids and oligonucleotides used in this study are listed in Table S3 and Table S4.

### RNA-sequencing

*M. tuberculosis* strains were grown in 25 ml of Sauton’s medium without detergent to OD_600_ ~ 0.5. Then, cultures were harvested by centrifugation and the resulting pellets resuspended in 1 mL TRIzol Reagent (ThermoFisher) and stored at −80°C until further processing. Bacteria were disrupted by bead-beating and total RNA was isolated as previously described (44). Two independent cultures for each strain were used for this experiment. Total RNA concentration was measured using the Qubit RNA HS Assay Kit. Library preparation, Illumina high-throughput sequencing and analysis were performed as described previously (36). The RNA-seq data were deposited at the Gene Expression Omnibus (GEO) database (https://www.ncbi.nlm.nih.gov/geo/query/acc.cgi?acc=GSE119582) under the accession number GSE119582.

### *In vitro* growth curves

All strains were diluted to an initial OD_600_ of 0.05 in Middlebrook 7H9 medium, and the OD_600_ recorded at different time points over a period of 7 days. Data from three independent experiments were used for growth representation.

### Antibiotic susceptibility

The MIC values were determined by using the resazurin-based microdilution assay (REMA) (28) as previously described (45). MIC values were determined by nonlinear fitting of the data to the Gompertz equation using GraphPad Prism.

### Macrophage survival

Human monocytic THP-1 cells were grown in RPMI medium supplemented with 10% FBS and 1% sodium pyruvate. Monocytes at 2×10^6^ cell/ml were differentiated into macrophages using phorbol-12-myristate-13-acetate (PMA) at a final concentration of 4 nM and subsequently pipetted at 10^5^ cell/well into a 96 well-plate. The Plate was sealed and incubated at 37°C with 5% CO_2_ for 18h. Medium containing PMA was removed and replaced with new RPMI medium. Bacteria were grown to OD_600_ 0.4-0.8, washed, resuspended in 7H9 broth to an OD_600_ of 1 (3 × 10^8^ bacteria/ml) and diluted in RPMI medium at a concentration of 10^7^ bacteria/ml. THP-1 cells were then infected at a multiplicity of infection (MOI) of 5 and incubated for two or three days at 37°C with 5% CO_2_. Afterwards, PrestoBlue Assay (Life Technologies) was used to evaluate cell viability according to the manufacturer’s instructions. Fluorescence was measured using a Tecan M200 instrument (excitation/emission wavelength of 560/590 nm). Fluorescence units of three biological replicates was analyzed in GraphPad Prism using the one-way ANOVA followed by Tukey’s multiple comparison test.

### Bacterial uptake by THP-1 macrophages

THP-1 macrophages were infected with *M. tuberculosis* strains at MOI of 5 and incubated at 37°C with 5% CO_2_ in the presence of RPMI. Four hours post-infection, extracellular bacteria were removed by washing twice with warm PBS and macrophages lysed with 0.1% TritonX-100 in PBS. Lysates were diluted, plated on 7H10 medium and incubated for three weeks before the CFU were determined. Results from three independent replicates are represented.

### Protein preparation, secretion analysis and immunodetection

Culture filtrates and cell lysates were prepared and immunoblotted as described previously (35) (16). GroEL2 was used as a loading control for cell lysates and Ag85B, an ESX-1 independent secreted protein, as a loading control for culture filtrate. The following reagents were purchased from Abcam: Monoclonal Anti-ESAT6 (Ab26246), polyclonal Anti-CFP10 (Ab45074) and polyclonal Anti-Ag85B (ab43019). Polyclonal Anti-EspB antibodies were produced by Dr. Ida Rosenkrands (Statens Serum Institut, Copenhagen, Denmark). The following reagent was obtained through BEI Resources, NIAID, NIH: Monoclonal Anti-*M. tuberculosis* GroEL2 (gene Rv0440), clone IT-56 (CBA1) (produced in vitro), NR-13655. Monoclonal Anti-HA tag antibodies conjugated to Horseradish Peroxidase (HRP) were purchased from Cell Signaling (#2999). Polyclonal Anti-Rv3852 antibody was produced by Eurogentec (30).

### Mass spectrometry-based analysis

Proteins for secretion analysis (culture filtrates) were prepared as described above. To prepare the cell lysate, the cell pellet was washed once with PBS, resuspended in 100mM Tris pH 8, 2% SDS buffer with Roche protease inhibitor cocktail tablets, disrupted by sonication for 15 min at 4°, clarified by centrifugation and heat-inactivated at 100°C for 1h. Total protein concentration in all preparations was determined using the BCA assays. Mass spectrometry analysis was performed as described previously (36). Significant hits were determined by a volcano plot-based strategy, combining t test *p*-values with ratio information (46). Significance curves in the volcano plot corresponding to a SO value of 0.5 and a permutation-based False Discovery Rate (FDR) of 0.05 were determined by a permutation-based method. The FDR was determined after 250 random permutations of the data allowing separation of biologically significant hits from those randomly obtained (46, 47). The mass spectrometry proteomics data have been deposited to the ProteomeXchange Consortium via the PRIDE (48) partner repository (https://www.ebi.ac.uk/pride/archive/simpleSearch?q=&submit=Search) with the dataset identifier PXD012584.

### Cell fractionation

*M. tuberculosis* Δ*eccE*_1_/*eccE*_1_*HA* mutant was grown in 100 ml of Sauton’s medium without Tween-80 for 4 days with a starting OD_600_ 0.5. Bacteria were harvested by centrifugation and the fractions were collected as described (30).

### Fluorescence microscopy

Bacteria expressing mNeon were grown in 7H9 broth to log phase (OD_600_ 0.4-0.8) then mounted on a microscope slide with 1.5% agarose pads and directly examined with an Olympus IX81 microscope under a 100x objective. Images were analyzed using ImageJ and normalized for brightness, contrast and resolution.

## Acknowledgments

The research leading to these results has received funding from the Swiss National Science Foundation under grant 31003A-162641 to STC. The funders had no role in study design, data collection and interpretation, or the decision to submit the work for publication.

We thank A. Benjak for the analysis of high-throughput sequencing data, C. Avanzi for library preparation, the Proteomics Core Facility at EPFL for mass spectrometry experiments, the Lausanne Genomic Technologies Facility at the University of Lausanne for high-throughput sequencing analyses and YW Chang for providing valuable advice.

## Supporting Information Captions

**FIG S1. EccE_1_ and MycP_1_ are not required for *in vitro* growth.**

Growth curves on synthetic medium (7H9 + ADC). The mutants and the WT strain display similar growth rates.

**FIG S2. Bacterial uptake by THP-1 macrophage.**

CFU enumeration of intracellular bacteria at 4h post-infection. THP-1 cells were infected at MOI of 5. Data are represented as the mean and SD of three independent replicates.

**FIG S3. EccE_1_-mNeon M. tuberculosis localises at polar regions only in the presence of all ESX-1 proteins.**

**A**. Δ*mycP*_1_-*eccE*_1_/*MycP*_1_-*eccE*_1_-mNeon cells. Representative image of three independent cultures demonstrating that 100% of cells expressing EccE_1_-mNeon displayed a bi-polar fluorescent pattern. **B.** ΔΔRD1/*mycP*_*1*_-*eccE*_*1*_-mNeon cells. Representative image of three independent cultures proving that EccE_1_-mNeon needs other ESX-1 protein to form a stable complex at the poles of the bacilli.

**FIG S4. Secretion of EspB in H37Rv and Erdman strains.**

Immunoblot of CF fractions. H37Rv mutants: ΔespB (knockout of espB), ΔmycP_*1*_-eccE_*1*_/*eccE*_*1*_ (knockout of mycP_1_), and Δ*mycP*_*1*_-*eccE*_*1*_/*mycP*_*1*_-*eccE*_*1*_. Erdman mutant: *pe35*::tn. Δ*espB* and *pe35*::Tn were used as negative controls as neither of these mutants secrete EspB. Ag85B was used as a loading control.

**Table S1.** Relative expression levels by RNA-sequencing.

**Table S2.** EccE_1_ and MycP_1_ are not involved in susceptibility to antibiotics targeting the cell wall.

**Table S3.** Strains and Plasmids used in this study.

**Table S4.** Oligonucleotides used in this study.

**Table S5.** Secretome of the mutants compared to that of the WT strain.

**Table S6.** Intracellular proteome of the mutants compared to that of the WT strain.

